# On the use of facial and neck cooling to improve indoor occupant thermal comfort in warm conditions

**DOI:** 10.1101/2021.10.24.465522

**Authors:** Bin Yang, Tze-Huan Lei, Faming Wang, Pengfei Yang

## Abstract

Face and neck cooling has been found effective to improve thermal comfort during exercise in the heat despite the surface area of human face and neck regions accounts for only 5.5% of the entire body. Presently, very limited work in the literature has been reported on face and neck cooling to improve indoor thermal comfort. In this work, two energy-efficient wearable face and neck cooling fans were used to enhance occupants’ thermal comfort in two warm indoor conditions (30 & 32 °C). Local skin temperatures and perceptual responses while using those two wearable cooling fans were examined and compared. Results showed that both cooling fans could largely reduce local skin temperatures at the forehead, face and neck regions up to 2.1 °C. Local thermal sensation votes at the face and neck were decreased by 0.82-1.21 scale unit at two studied temperatures. Overall TSVs dropped by 1.03-1.14 and 1.34-1.66 scale unit at 30 and 32 °C temperatures, respectively. Both cooling fans could extend the acceptable HVAC temperature setpoint to 32.0 °C, resulting in an average energy saving of 45.7% as compared to the baseline HVAC setpoint of 24.5 °C. Further, the free-control cooling mode is recommended to occupants for further improving thermal comfort while using those two types of wearable cooling fans indoors. Lastly, it is concluded that those two wearable cooling fans could greatly improve thermal comfort and save HVAC energy despite some issues on dry eyes and dry lips associated with those wearable cooling fans were noted.

## 1. Introduction

Personal thermal management (PTM) has received tremendous attention over the past years as it helps save building energy and improve individual occupant thermal comfort [1–3]. In general, a personal thermal management system (PTMS) creates an ideal near-body thermal envelop so that individuals’ thermal comfort could be improved. Besides, personal thermal management systems consume very little energy compared to traditional HVAC (heating, ventilation, and air-conditioning) systems [4].

PTMS can be categorized into two types: wearable PTMS and non-wearable PTMS. Non-wearable PTMS may include such old-fashion devices as task ambient conditioning systems, personal comfort devices, and personalized ventilation systems [5–9]. The use of non-wearable PTMS to provide individual occupants thermal comfort has been extensively examined over the past 3 decades [5–9]. Documented investigations on non-wearable PTMS have eloquently elucidated that the use of non-wearable PTMS could further improve individual thermal comfort in both non-air-conditioned and air-conditioned indoor environments [3]. A recent review article by Yang et al. [3] pinpointed that individual thermal comfort could be further improved if the intensified conditioning of personal micro-environment is moved closer to the body. Hence, it is anticipated that wearable PTMS may further improve individual thermal comfort while consuming minimal energy or even no energy. Presently, wearable PTMS may be further classified into two groups: PTMS incorporated with cooling/heating modules and clothing made of specially designed materials and/or with a unique fabric layer structure. Ke et al. [10] examined the performance of nanoporous polyethylene (nanoPE) passive cooling clothing on the improvement of occupants’ indoor thermal comfort. It was found that the nanoPE passive cooling clothing could extend indoor acceptable air-conditioning setpoint temperature to 27.0 °C and thereby, saving 9-15% cooling energy. Ma et al. [11] numerically analyzed the energy saving performance of novel radiative cooling PTMS. Results indicated that personal radiative cooling textiles (with air gap size of 5 mm) could save 4.6-12.8% energy in worldwide locations. At present, very limited literatures reported the use of wearable PTMS incorporated with cooling units to improve occupants’ thermal comfort both locally and at the whole-body level. Song et al. [12] explored the effect of hybrid personal cooling clothing on enhancement of thermal comfort of office workers in a hot indoor condition (34 °C, 65%RH [relative humidity]). Results demonstrated that hybrid cooling clothing could effectively improve local- and whole-body thermal comfort. Udayraj et al. [13] assessed and compared the performance of a conventional desk fan and air ventilation clothing incorporated with micro fans in three warm indoor conditions (28, 30 and 32 °C). Results indicated that both systems showed similar perceptual response and skin temperature at all three air temperatures. Air ventilation clothing could save 7-8% energy as compared to the conventional desk fan. Wang et al. [14] evaluated comfort of a thermally dynamic wearable thermoelectric wrist band. This device could improve overall thermal sensation, comfort and pleasantness 0.5-1 scale unit. However, the above results were obtained in thermal neutral conditions (<26.0 °C) and hence, the findings might not be translated into warmer indoor conditions. On the other hand, air ventilation clothing reported in aforementioned studies had some practicality limitations. For instance, such clothing became rather bulky during operation and also, some hygienic issues related to contaminated air due to sweating/body scent might not be avoided. Therefore, there is a need to look for better wearable personal cooling systems to improve local body cooling while working in indoor environments.

Local body thermal sensitivity to heat stress environments should be considered while applying wearable PTMS for improving thermal comfort. Arens et al. [15] pointed out that the head is insensitive to cold environments but sensitive to warm environments. Literatures [5, 16] showed that face cooling could improve occupants’ thermal acceptability and thereby shifting the acceptable upper boundary of indoor temperature. Cotter and Taylor [17] found that the head/face and neck regions have much greater alliesthesial responses than the rest of the body. Though the cooling area of face and neck regions are relatively small, remarkable effectiveness on the thermal comfort improvement of human subjects could be anticipated because of high sensitivity at the face and neck areas. Thus, it was hypothesized that cooling of the head/face and neck region using wearable PTMS could bring a pronounced thermal comfort improvement on occupants while performing office work in warm indoor conditions.

It is also worth mentioning that current literatures concerning PTMS often adopted the fixed-power cooling module whilst neglecting to investigate the role of individual behaviour response on PTMS [10, 13]. It is well established that individual occupant may have their own preferred cooling temperatures. Given this fact, the individual free-control cooling module may be more effective than the fixed-power cooling method on thermal comfort improvement for indoor occupants while working in warm environments. Contrary to the above expectation, Boerstra et al. [18] discovered that task performance was better when participants had no control as compared to free control of personal desk fans. Hence, further investigations are still required to address and compare the impacts of free-control and fixed-power (i.e., no control) on occupants’ thermal comfort.

In this work, two types of highly energy-efficient (power consumption ≤ 4 W) wearable cooling fans (face cooling fan & neck cooling fan) were chosen to examine their actual performance on the enhancement of occupants’ thermal comfort while doing office work in two warm indoor conditions. The impact of those two wearable cooling systems on occupants’ overall and local physiological and perceptual responses was extensively investigated. Furthermore, the effectiveness of face and neck cooling on thermal comfort was compared and discussed. Lastly, two cooling control modes including fixed-power and free-control modes were chosen to further examine how did personal control mode affect occupants’ thermophysiological and perceptual responses. This work may serve as a useful guide for practitioners on how to wisely operate wearable personal cooling systems for the enhancement of individual thermal comfort in warm conditions.

## 2. Methods

### 2.1 Participants

Assuming an effect size of 0.65, α=0.05, and power of 0.8, eleven participants could provide enough power to register a statistical difference of a similar magnitude (G*Power Version 3.1.9.6, Heinrich-Heine-Universität Düsseldorf, Düsseldorf, Germany). Hence, sixteen young college students (8 males and 8 females) participated in this work. The physical characteristics of 16 participants are shown in Table 1. All participants were physically healthy and had no history of heat illnesses, pulmonary, or cardiovascular diseases. They were advised not to drink tea, coffee, alcohol and perform any intensive activity at least a day before each trial. Prior to participation, participants were fully briefed of the purpose and details associated with this study. A written informed consent was acquired thereafter. Participants might quit this study at any time without penalty. They received an honorarium after completed all trials.

**Table 1.** Physical characteristics of participants.

| Gender | Age<br>(yr) | Height<br>(m) | Weight<br>(kg) | Body mass index<br>(kg/m<sup>2</sup>) | Body surface<br>area (m<sup>2</sup>) |
| --- | --- | --- | --- | --- | --- |
| Males | 23.8±1.7 | 1.76±0.04 | 67.13±8.29 | 21.70±1.85 | 1.82±0.13 |
| Females | 22.6±2.0 | 1.64±0.05 | 55.13±3.87 | 20.62±1.71 | 1.59±0.07 |
| Overall | 23.1±2.3 | 1.70±0.08 | 66.13±8.80 | 21.16±1.81 | 1.70±0.15 |

Note: data are presented as mean±SD (standard deviation).

### 2.2 Face and neck cooling fans

In order to examine the actual performance of energy-efficient wearable cooling fans on the improvement of occupants’ thermal comfort in warm indoor conditions, two commercially-available wearable U-shaped cooling fans were selected: a face cooling fan and a neck cooling fan. The face cooling fan (Gusgu, Shenzhen Gushang Digital Co., Ltd., Shenzhen, China) uses two brushless 360° rotating axial fans to generate airflow and can be worn around the neck. The two axial fans have a diameter of 7.5 cm and they can be operated at three adjustable speed levels. This wearable face cooling fan has a bult-in rechargeable 2000 mA·h (7.4 Wh, voltage: 3.7 V) lithium battery and can be recharged via a USB cable. The cooling duration lasts for 3-10 hours depends on speed levels (air speed ranged from 2.20-4.00 m/s at levels 1-3 [which was measured using an anemometer at a distance of 10 cm]). The total power and total weight of the face cooling fan is 4.0 W and 220 g, respectively. For the wearable neck cooling fan (Gusgu WT-F41, Shenzhen Gushang Digital Co., Ltd., Shenzhen, China), two 5-cm (diameter) brushless centrifugal fans were used. The air generated by the two centrifugal fans blows out from 76 tiny vents located along the U-shape air ducts. This type of neck cooling fan is powered by a 2400 mA·h (8.9 W·h, voltage: 3.7 V) rechargeable lithium battery and the cooling duration last for 3-16 hours depends on the fan speed. Similar to the wearable face cooling fan, this neck cooling fan can also be operated at three levels of air speed (averaged air speed at the outlets was 1.15-3.25 m/s at speed levels 1-3). The total power and total weight of the neck cooling fan is 3.7 W and 260 g, respectively.

### 2.3 Test protocol & procedure

Each participant completed 12 trials [2 temperature x 3 cooling options x 3 cooling modes] at two levels of air temperature (i.e., 30 and 32 °C), with three cooling options (i.e., CON [no cooling], FC [face cooling using the face cooling fan] and NC [neck cooling using the neck cooling fan]), and two cooling control modes (i.e., fixed power at the speed level 2 [fixed], and freely control the fan speed [free-control]). The selection of the speed level 2 in the fixed-power mode was based on the participants’ feedback from a pilot trial regarding the most often used fanning speed during practice. Hence, there are 192 test scenarios in total. All trials were randomized and counter-balanced and performed at the same time of the day.

Upon arrival at the laboratory, participants rested in armchairs for 20-30 min. Thereafter they were instrumented with skin temperature sensors. Local skin temperatures at the forehead, face and the neck were measured using wireless skin temperature loggers (iButton DS1922L, Maxim Integrated, San Jose, CA; resolution: 0.0625, accuracy: ±0.5 °C). Participants were dressed in underwear (briefs, panties, bra [for females]), long trousers, a short sleeve t-shirt (100% polyester), socks, a pair of shoes (estimated total clothing thermal insulation is 0.57 clo). Afterwards, they entered the climatic chamber (dimension: 3800×3800×2600 cm^3^) and were seated around a table. Participants could choose either reading books or working with computers during the entire trials (estimated metabolic rate was 1.0 met).

In all trials, occupants’ perceptual responses including overall and local-body thermal sensation votes (TSVs), overall thermal comfort votes (TCVs), dry eyes and lips were surveyed throughout the entire trials at 10 min intervals (detail of perceptual rating scales is addressed in section 2.5). The total duration of each exposure trial is 50 min. The air temperature, relative humidity (RH), air speed and the carbon dioxide concentration inside the climatic chamber were recorded at an interval of 1 min. Local skin temperatures at the forehead, face and the neck were collected every 1 min as well. Table 2 illustrates the equipment adopted for recording environmental parameters in the chamber.

**Table 2.** Details of measurement equipment used in this study.

| Parameters | Instrument (type and manufacturer) | Accuracy |
| --- | --- | --- |
| Air temperature | HOBO U12–012 (Onset Corp., Bourne, USA) | $\pm 0.35$ °C |
| Relative humidity | HOBO U12–012 (Onset Corp., Bourne, USA) | $\pm 2.5\%$ |
| Air speed | Swema 03 anemometers (Swema AB, Farsta, Sweden) | $\pm 0.05$ m/s |
| CO<sub>2</sub> concentration | RTR-576 (T&D Corporation, Nagano, Japan) | $\pm 50$ ppm |

Note: ppm, parts per million.

### 2.4 Test conditions

Two indoor air temperatures were chosen for this study, i.e., 30 and 32 °C. The operative temperature inside the chamber was assumed to be equal to the air temperature because the wall temperature was maintained at the same temperature as the ambient air. The indoor relative humidity was maintained at 50±5% and the air speed was 0.1 m/s. The partial water vapor pressure in the chamber was 2.16 and 2.42 kPa at 30 and 32 °C temperatures, respectively. Based on the CBE thermal comfort tool [19], the PMV (predicted mean vote) was +1.55 and +2.29 at air temperatures of 30 and 32 °C, respectively.

### 2.5 Perceptual response questionnaire

E-questionnaire was adopted to collect overall and local-body perceptual responses of occupants. Overall perceptual responses included the ASHRAE 7-point thermal sensation vote (TSV) [20], thermal comfort vote (TCV), and ratings of dry eyes and dry lips [10]. TCV scale ranged from ‘Very uncomfortable’ (−3), to ‘Uncomfortable’ (−2), to ‘Slightly uncomfortable’ (−1), to ‘Neutral’ (0), to ‘Slightly comfortable’ (+1), to ‘Comfortable’ (+2), and to ‘Very comfortable’ (+3). Ratings of dry eyes and dry lips ranged from ‘Dry’ (−2), to ‘Slightly dry’ (−1), to ‘Neutral’ (0), to ‘Slightly wet’ (+1), to ‘Wet’ (+2). All rating scales are continuous except rating scales of dry eyes and dry lips (discrete scales). The questionnaire appeared automatically on the occupants’ computer screen every 10 min throughout the entire trials, and data were stored in the computer. Participants spent about 60 seconds to complete all survey questions.

### 2.6 Statistical analysis

Steady-state data (i.e., the last 20 min of each trial) were analyzed and reported. Data are reported as mean±SD (standard deviation) and were evaluated for normality with the Shapiro-Wilk test. Violations of Mauchly’s test of sphericity were adjusted using Greenhouse Geisser adjustments. Three-way repeated-measures ANOVA was performed to examine whether independent variables (i.e., cooling conditions [CON, FC & NC], air temperatures [30 & 32 °C] and cooling modes [fixed & free-control]) significantly affected such dependent variables as local skin temperatures at the forehead, face and the neck, as well as overall and local perceptual responses. If a significant difference was detected, Paired Samples t-tests were performed to examine which pairs of test scenarios had significant differences. All statistical analyses were carried out using SPSS Statistics Version 26.0 (IBM, Chicago, IL). The significance level was set as *p*<0.05.

## 3. Results

### 3.1 Local skin temperatures at forehead, face and neck

Local mean skin temperatures at the forehead, face and neck in the 12 studied test scenarios are presented in Figures 1(a), (b) and (c), respectively. In both two studied air temperatures (i.e., 30 and 32 °C), the use of wearable face and neck cooling fans significantly reduced local skin temperatures at the forehead, face and the neck (*p*<0.001). The forehead temperature decreased by 0.3-1.0 °C in FC and NC as compared to CON. At 30 °C, free-control of the fans raised local skin temperatures at the forehead and the mean forehead temperature was 34.8 and 35.4 in FC30(free-control) and NC30(free-control), respectively. In contrast, the free-control mode reduced the forehead skin temperature by 0.2 °C compared to the fixed power mode in FC at 32 °C. No significant difference in the mean forehead temperature was observed between NC32(fixed) and NC32(free-control).

**Figure 1.**
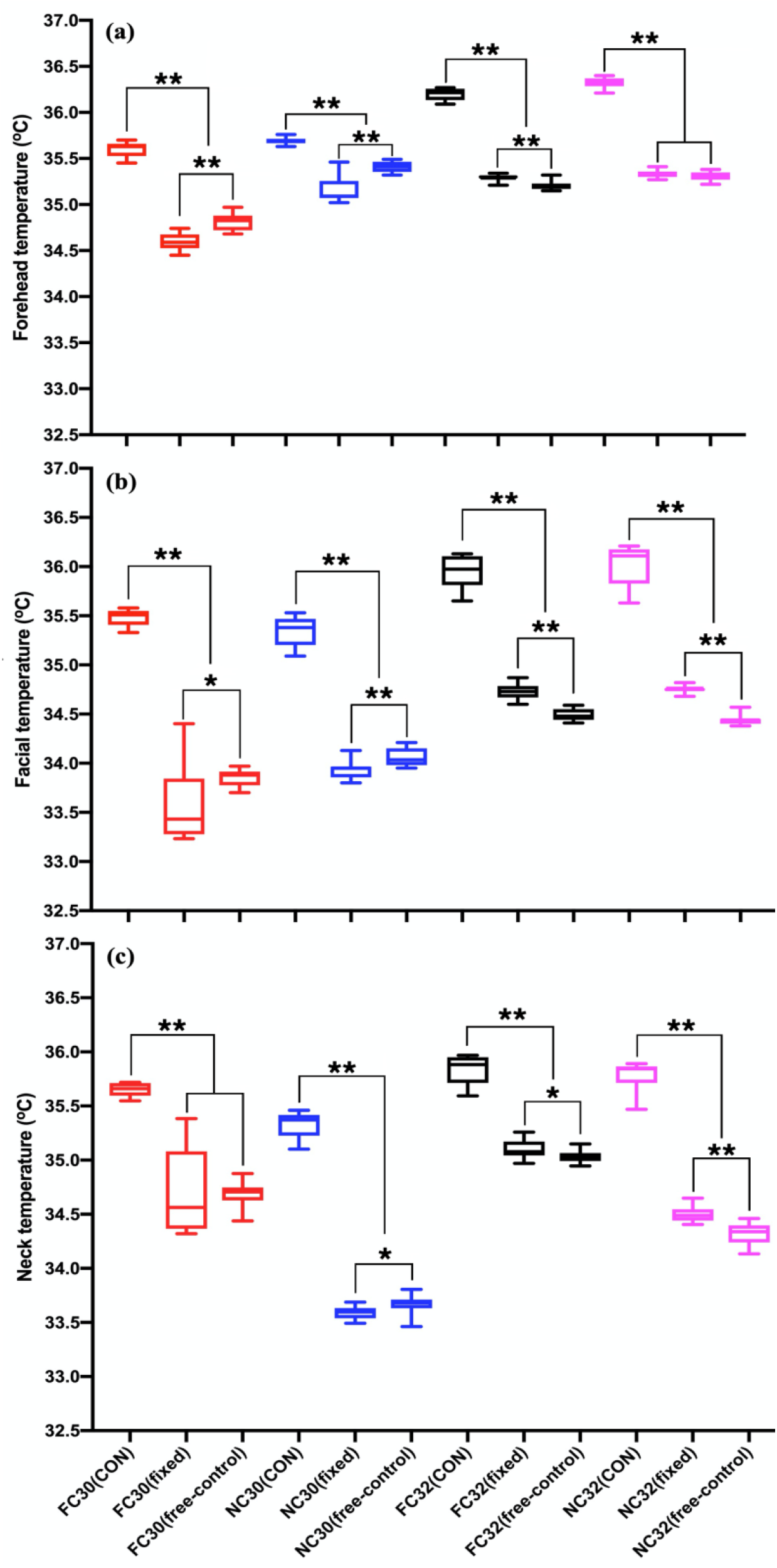
Local mean skin temperatures at the (a) forehead, (b) face and (c) neck of 12 studied test scenarios. *, *p*<0.05; **, *p*<0.001.

With regard to the mean face temperature, it was decreased by 1.4-1.9 °C at 30 °C in FC and NC as compared to no cooling. Similarly, the mean face temperature decreased by 1.2-1.6 °C at 32 °C temperature when the two wearable cooling fans were applied. Furthermore, it was observed that the face temperature increased by 0.2-0.3 °C in the free-control mode as compared to the fixed-power mode at 30 °C. In contrast, the face temperature decreased by 0.2-0.4 °C in the free-control mode as compared to the fixed-power mode at 32 °C. The mean face skin temperature was 34.7, 34.5, 34.8 and 34.4 °C in FC32(fixed), FC32(free-control), NC32(fixed) and NC32(free-control), respectively. Further, it was noted that local face temperature was significantly lower in face cooling fan scenarios than the neck cooling fan scenarios.

As for the neck skin temperature, the use of face cooling fan could only decrease the neck temperature by 0.7-1.0 °C whereas the neck cooling fan could reduce the local neck skin temperature by 1.3-2.1 °C. Significant differences were registered in the mean neck temperature between the fixed-power mode and the free-control mode in NC30 (*p*<0.001), FC32 (*p*<0.05) and NC32 (*p*<0.001). The mean neck skin temperature was 34.7 and 34.7 °C in FC30(fixed) and FC30(free-control), respectively. It was 33.6 and 33.7 °C in NC30(fixed) and NC30(free-control), respectively. At 32 °C, the mean neck skin temperature was 35.1, 35.0, 34.5 and 34.3 °C in FC32(fixed), FC32(free-control), NC32(fixed) and NC32(free-control), respectively.

### 3.2 Overall thermal sensation

Overall thermal sensation votes (TSVs) of the 12 studied test scenarios are shown in Figure 2. The observed overall TSVs were +1.73 (close to ‘Warm’), +1.66 (close to ‘Warm’), +2.44 (in between ‘Warm’ and ‘Hot’) and +2.40 (in between ‘Warm’ and ‘Hot’) in FC30(CON), NC30(CON), FC32(CON), NC32(CON), respectively. The use of two energy-efficient cooling fans significantly improved overall TSVs in all test scenarios (*p*<0.001). Overall TSVs decreased to +0.59 to +0.64 (in between ‘Neutral’ and ‘Slightly Warm’) when face and neck cooling fans were used at 30 °C. In contrast, it decreased to +0.74 to +1.10 (close to ‘Slightly Warm’) if face and neck cooling fans were applied at 32 °C. Further, the free-control mode did not significantly affect the overall TSVs as compared to the fix-power mode in all test scenarios (*p*>0.05) except NC32 scenarios (*p*<0.05).

**Figure 2.**
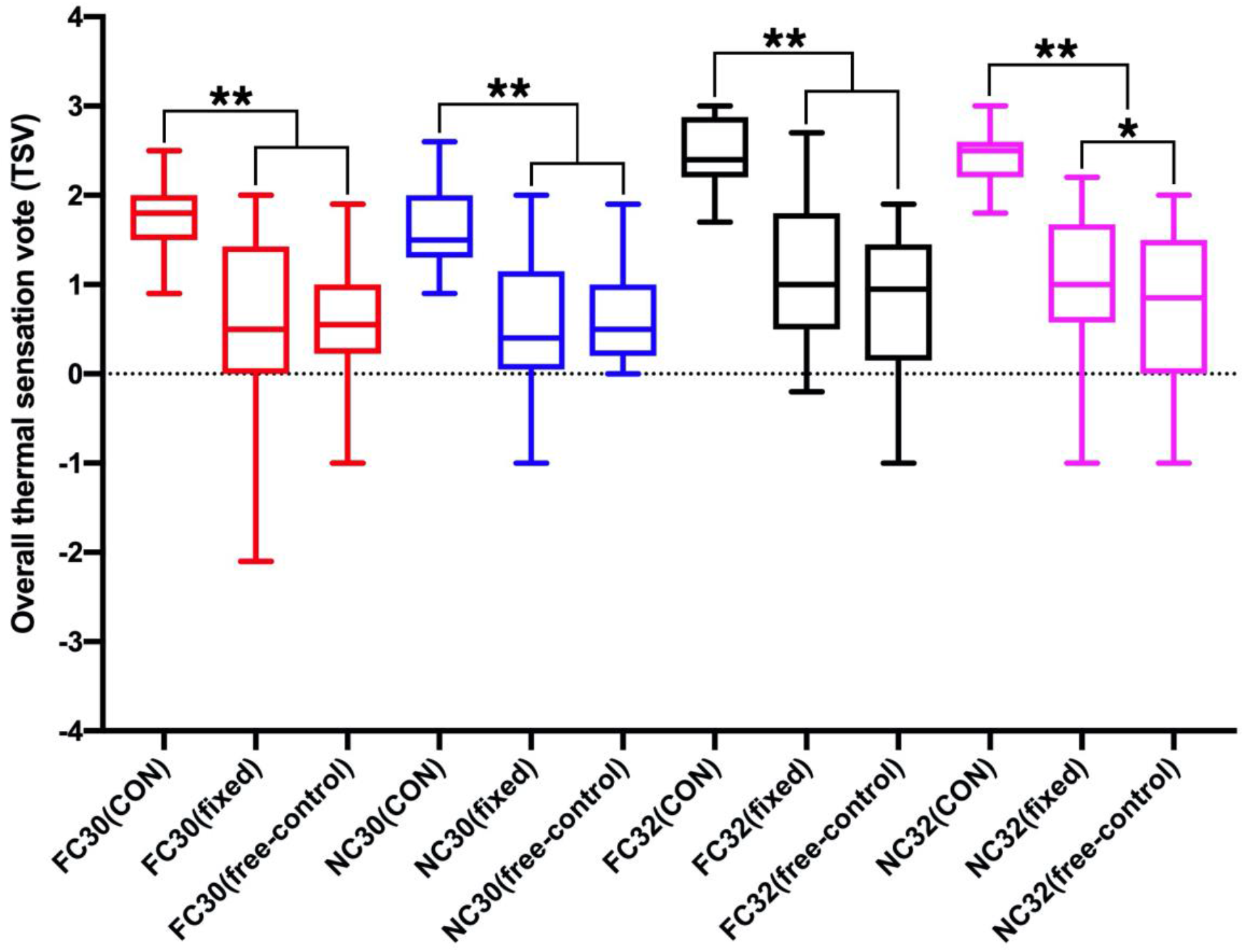
Overall thermal sensation votes (TSVs). *, *p*<0.05; **, *p*<0.001.

### 3.3 Overall thermal comfort

Overall thermal comfort votes (TCVs) of the 12 studied test scenarios are displayed in Figure 3. Overall TCVs were −0.85 and −0.65 (close to ‘Slightly Uncomfortable’) when no cooling was adopted at 30 °C. In contrast, overall TCVs were −1.12 and −1.13 (close to ‘Slightly uncomfortable’) at 32 °C in CON. The use of two wearable cooling fans significantly improved overall TCVs compared to no cooling fan (*p*<0.05 or *p*<0.001). Overall TCVs were improved by 0.30-0.96 scale unit when the two wearable cooling fans were used at both air temperatures. It is interesting to observe that the free-control mode could significantly improve overall TCVs as compared to the fixed-power mode at 30 °C (p<0.05). Overall TCVs in FC30(free-control) and NC30(free-control) were +0.11 and −0.07 (close to ‘Neutral’), respectively. Conversely, no significant differences in overall TSVs were discovered between the fixed-power and free-control modes with both two cooling fans at 32 °C (*p*>0.05).

**Figure 3.**
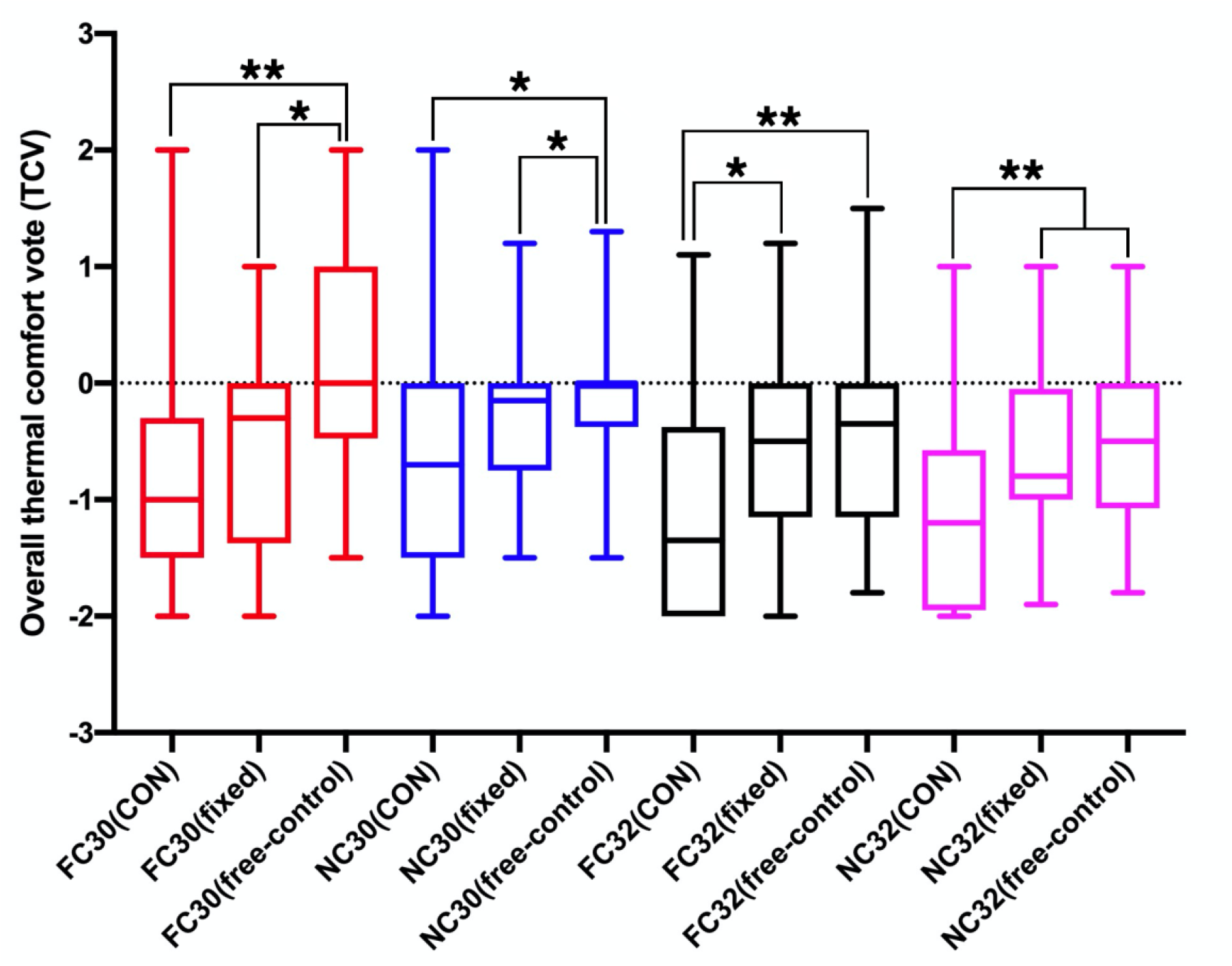
Overall thermal comfort votes (TCVs). *, *p*<0.05; **, *p*<0.001.

### 3.4 Local thermal sensation at face and neck

Local thermal sensation votes (TSVs) at the face and the neck areas are illustrated in Figures 4(a) and (b), respectively. Compared to no cooling, the two wearable cooling fans significantly improved local thermal sensation at the face area as well as the neck (*p*<0.001). Local TSVs at the face area decreased by 0.78-1.20 scale unit by the cooling fans at 30 °C, whereas it was reduced by 0.98-1.21 scale unit at the air temperature of 32 °C. During the cooling period, all local TSVs at the face area were maintained at below +0.61 (in between ‘Neutral’ and ‘Slightly warm’) at both 30 and 32 °C. In addition, the face cooling fan showed greater improvement on local face TSVs than the neck cooling fan. With regard to the control mode, the free-control mode showed significantly higher local TSVs at the face than the fix-power mode in NC30 scenarios (*p*<0.05).

**Figure 4.**
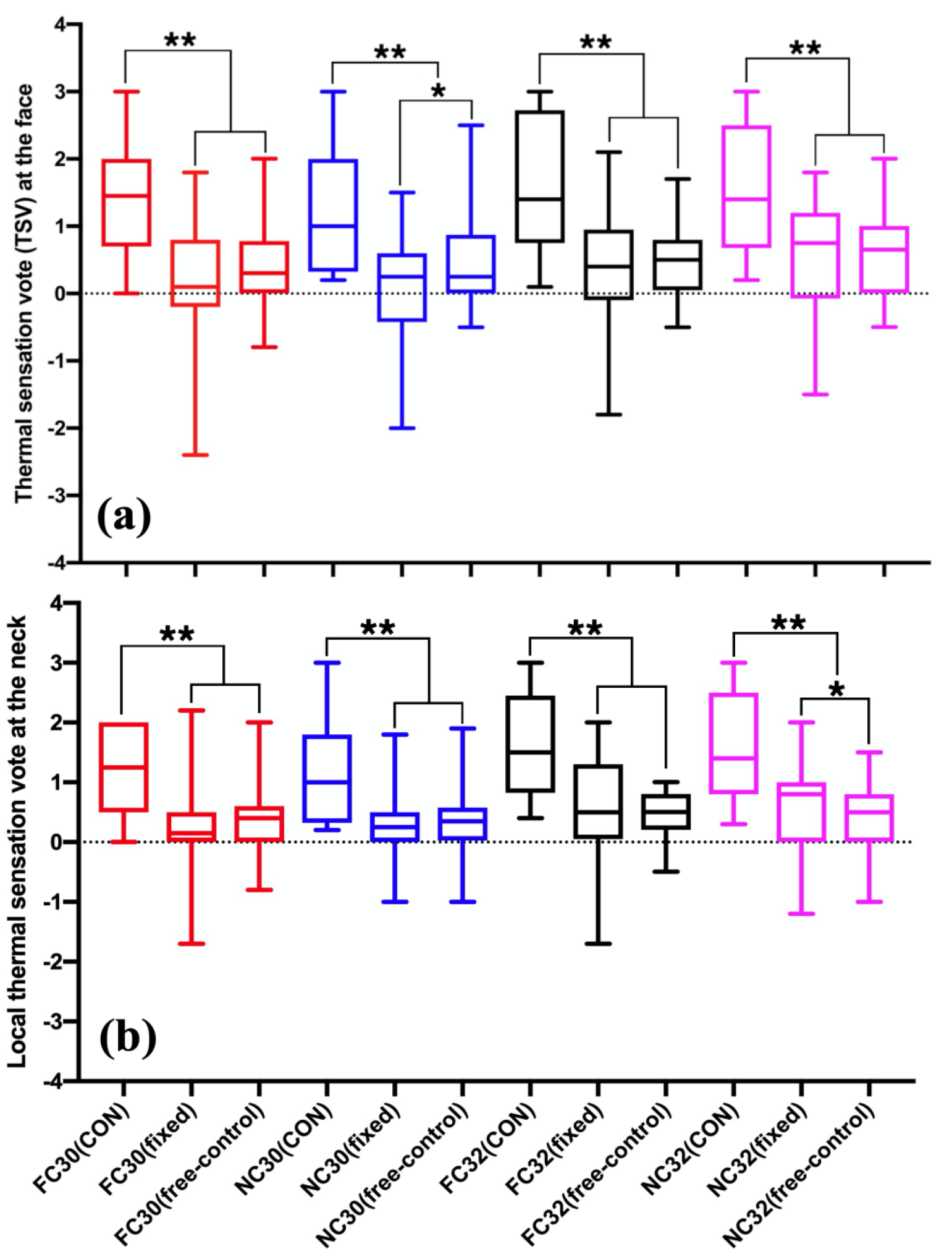
Local thermal sensation votes at (a) the face area and (b) the neck. *, *p*<0.05; **, *p*<0.001.

As for mean local TSVs at the neck, local TSVs at the neck were in between +1.19 and +1.62 (close to ‘Slightly warm’) when no cooling was applied [see Figure 4(b)]. Both cooling fans significantly reduced local TSVs at the neck area by over 0.82 scale unit. Local TSVs at the neck were +0.29, +0.43, +0.24 and +0.36 (all values were close to ‘Neutral’) in FC30(fixed), FC30(free-control), NC30(fixed) and NC30(free-control), respectively. Similarly, Local TSVs at the neck were +0.55, +0.44, +0.52 and +0.37 (in between ‘Neutral’ and ‘Slightly warm’) in FC32(fixed), FC32(free-control), NC32(fixed) and NC32(free-control), respectively. Hence, the neck cooling fan could induce greater improvement on local TSVs at the neck compared to the face cooling fan. The free-control mode showed significantly lower TSVs at the neck compared to the fix-power mode at 32 °C only (*p*<0.05).

### 3.5 Dry eyes and dry lips

Ratings of dry eyes and dry lips in the 12 studied test scenarios are demonstrated in Figures 5 and 6, respectively. It can be seen from Figure 5 that the use of both two cooling fans raised the percentage of participant reported dry eyes (ratings of −2 and −1). For instance, only 12.5-25% of participants rated dry eyes when no cooling was used, whereas the percentage of participants rated dry eyes of −2 and −1 increased to 50-68.8% when the two wearable cooling fans were used. On the other hand, the face cooling caused up to 15.6% higher percentage of dry eyes compared to the neck cooling fan. The free-control mode could largely improve the dry eye issues by 18.8% compared to the fix-power mode with the face cooling fan at 30 °C. On the contrary, the free-control mode worsened the dry eyes by 6.25% with the face cooling fan at 32 °C compared to the fix-power mode.

**Figure 5.**
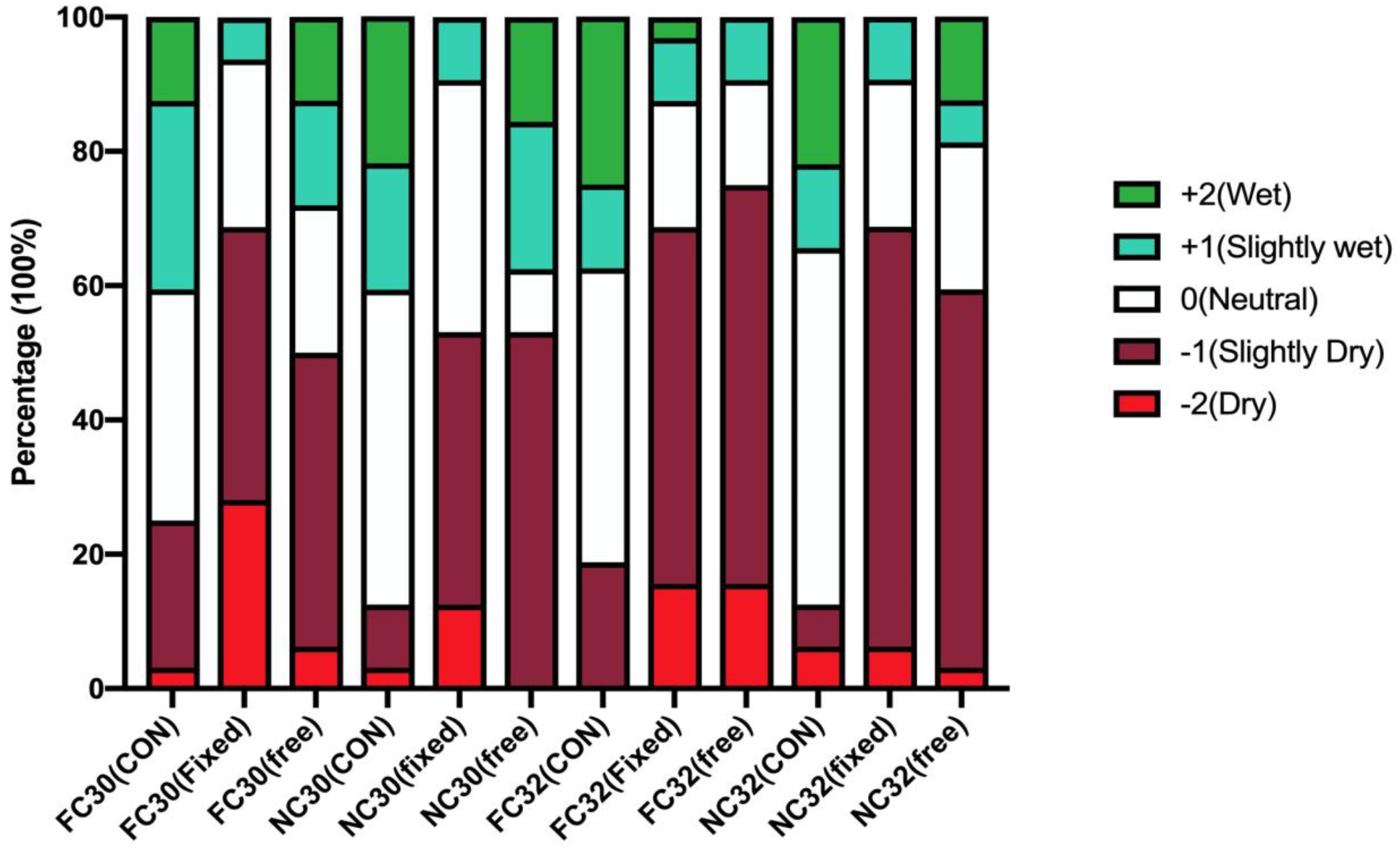
Ratings of dry eyes in the 12 studied test scenarios.

**Figure 6.**
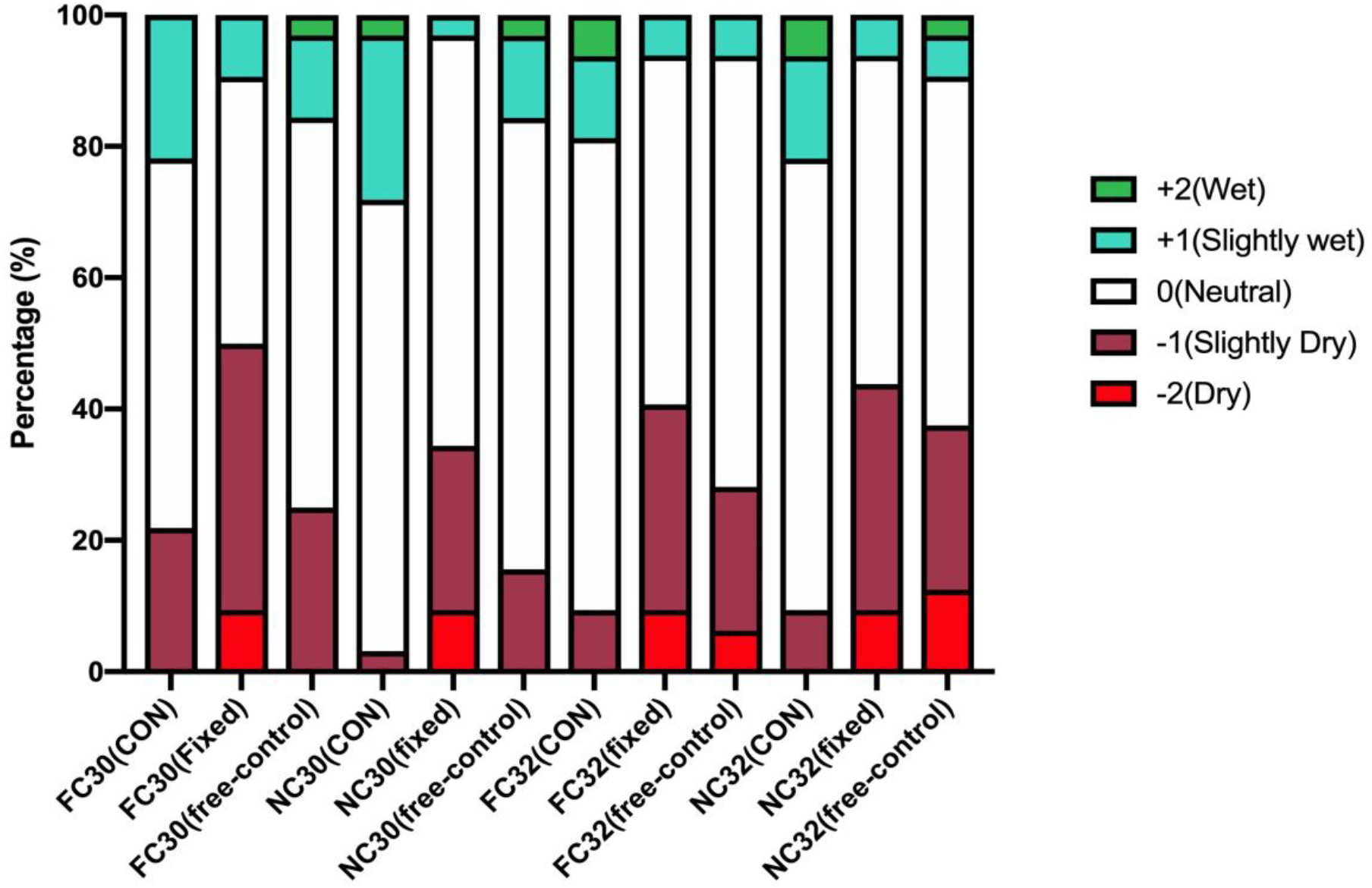
Ratings of dry lips in the 12 studied test scenarios.

As for the ratings of dry lips, the use of two wearable cooling fans greatly worsened the dry lips issue compared to no cooling (i.e., only less than 21.9% of participants reported dry lips issue [see Figure 6]). The percentage of dry lips raised from 9.4-21.9% to 15.6-50.0% when cooling fans were used. At the air temperature of 30 °C, the neck cooling fan showed 9.4-15.6% lower of dry lips than the face cooling fan. In all cooling fan cases, the free-control cooling mode greatly improved the dry lips issue by 6.3-18.8% as compared to the fixed-power cooling mode.

## 4. Discussion

The surface area of the human face and neck accounts for only 3.5% and 2% of the entire body surface area of an adult, respectively [21]. Face and neck cooling has been widely used to improve athletic performance in the heat [22–25]. In particular, neck cooling during exercise was found effective to improve exercise performance in the heat [23, 26, 27]. It has been well known that the activity intensity was pretty high during exercise and sports, still, the neck and face cooling functioned well. In indoor environments, occupants’ activity was much lower than athletes. Hence, face and neck cooling should be even more effective to improve thermal comfort of occupants compared to cooling athletes in the heat.

To the best of our knowledge, this is the first study to investigate wearable face and neck cooling on the improvement of thermal comfort of indoor occupants. Previous studies on electric fans [13, 28–32] have consistently shown electric fans could improve occupants’ thermal comfort in various warm indoor conditions. Nevertheless, such electric fans exposed the occupants to either whole-body convective cooling or upper-body cooling (including face and neck cooling). The performance of face and neck cooling on the enhancement of occupants’ thermal comfort in indoor conditions, however, has remained unknown. Results from this work have demonstrated that wearable face and neck cooling fans could greatly reduce local skin temperatures at the forehead, face as well as the neck region by up to 2.1 °C. It was also noted that face cooling fan could result in higher temperature reduction at the forehead and the face as compared to the neck cooling fan. Conversely, the neck cooling fan could cause higher skin temperature reduction at the neck as compared to the face cooling fan (see Figure 1). Further, local thermal sensation votes at the face have been improved by up to 1.21 scale unit at the two studied air temperatures (i.e., 30 and 32 °C). Similarly, the wearable face and neck cooling fans decreased the local thermal sensation at the neck by over 0.82 scale unit. Zhang and Zhao [33] examined the effect of face cooling (supplied by a personalized ventilation system) on human responses and found that the acceptable room temperature range could be raised from 26 to 30.5 °C while face cooling was provided. In this work, it seemed that the acceptable room temperature could further be raised to 32.0 °C while wearable face and neck cooling fans were used. Therefore, this could result in an average saving of 45.7% compared to the baseline HVAC setpoint of 24.5 °C [34].

Energy-efficient wearable face and neck cooling fans (power consumption ≤4 W) improved not only local thermal sensation at the face and the neck, but also significantly improved the overall thermal sensation as well as overall thermal comfort. Overall TSVs were +1.66 to +1.73 at 30 °C air temperature whereas TSVs were +2.40 to +2.44 at 32 °C. The PMV predicted by the CBE thermal comfort tool was +1.55 and +2.29 at those two indoor conditions (30 and 32 °C), respectively. It is evident that our observed TSVs were in good agreement with PMVs predicted by the CBE thermal comfort tool [19]. When the wearable face and neck cooling fans were used, overall TSVs reduced by 1.03-1.14 scale unit at 30 °C and they decreased by 1.34 to 1.66 scale unit at higher temperature of 32 °C. The observed overall TSVs were close to +0.5 (‘Slightly warm’) when face and neck cooling fans were used at 30 °C, which revealed that about 90% of participants showed thermal comfort satisfaction [35]. At 32 °C, the observed overall TSVs were close to +1.0 and hence, about 26% occupants were dissatisfied with the thermal condition (with cooling, PPD [Predicted Percentage of Dissatisfied]=88%). Thus, the use of face and neck cooling fans could result in 74% of occupants being satisfied with the thermal environment. Nevertheless, this number is slightly below the 80% occupant satisfaction rate defined by ASHRAE 55 [20]. This could be due to the dry eyes and lips issue caused by the use of face and neck cooling fans (see Figures 5–6). On the other hand, overall TCVs at 32 °C were all above −0.72 (see Figure 3), which denoted that the 32 °C temperature was still acceptable when using such energy-efficient wearable face and neck cooling fans.

The impact of cooling control mode on thermal comfort has also been examined in this work. At 30 °C, the fixed-power control mode caused overcooling in the face and neck regions, which was evidenced by lower overall and local TSVs as well as the lower TCVs as compared to the free control mode (see Figures 1–4). In contrast, at 32 °C, the fixed-power at the speed level 2 (corresponding wind speed: 2.18 m/s) was not able to bring sufficient cooling to the occupants. Hence, overall and local TSVs as well as overall TCVs were worsened with the fixed-power mode as compared to the free-control mode. The above findings verified that personal control (free-control mode) played a vital role in improving individual thermal comfort. This is consistent with previous work demonstrating that that individual control of personal comfort devices could increase an individual’s satisfaction with indoor conditions and energy efficiency [36–40].

Some limitations of this study should be acknowledged. First, only young college students were recruited, restricting our findings to populations with different ages and vulnerable groups. Second, local thermal comfort at the face and the neck was not examined and such details may provide useful information to explore its impact on overall thermal comfort. Next, only natural air cooling was studied and other wearable cooling options such as liquid cooling and evaporative cooling were not investigated. Future studies should be performed to evaluate the effectiveness of such energy-efficiency face and neck cooling fans on the elderly under various temperature conditions. Besides, various types of wearable cooling devices should be explored to look for the best performance wearable cooling devices for improving indoor occupants’ thermal comfort under higher temperatures.

## 5. Conclusions

Two energy-efficient wearable face and neck cooling fans were used to improve occupants’ thermal comfort in two warm indoor conditions. Human physiological and perceptual responses while using those two types of wearable cooling fans were examined and compared. Results showed that both wearable cooling fans could largely reduce local skin temperatures at the forehead, face as well as the neck regions up to 2.1 °C. Local thermal sensation votes at the face and the neck were decreased by 0.82-1.21 scale unit. Overall TSVs reduced by 1.03-1.14 scale unit at 30 °C and they decreased by 1.34-1.66 scale unit at 32 °C. Both cooling fans could extend the acceptable HVAC temperature setpoint to 32.0 °C, resulting in an average energy saving of 45.7% as compared to the baseline HVAC setpoint of 24.5 °C. Further, the free-control cooling mode is recommended to occupants for further improving thermal comfort while using those two types of wearable cooling fans indoors. It is ultimately concluded that the selected two types of wearable cooling fans could greatly improve thermal comfort and save HVAC energy despite some issues on dry eyes and lips were noted.

## Author contributions

Original idea: FW; Development and formulation of concept: BY, FW, THL & PY; Draft: YB, THL, & FW; Critical revision: THL, & FW.

## References

[1] J.K. Tong, X. Huang, S.V. Boriskina, J. Loomis, Y. Xu, G. Chen, Infrared-transparent visible-opaque fabrics for wearable personal thermal management, ACS Photon. 2 (2015) 769–778.

[2] Y. Peng, Y. Cui, Advanced textiles for personal thermal management and energy, Joule 4 (2020) 724–742.

[3] B. Yang, X. Ding, F. Wang, A. Li, A review of intensified conditioning of personal micro-environments: moving closer to the human body, Energ. Built Environ. 2 (2021) 260–270.

[4] H. Zhang, E. Arens, Y. Zhai, A review of the corrective power of personal comfort systems in non-neutral ambient environments, Build. Environ. 91 (2015) 15–41.

[5] A.K. Melikov, Personalized ventilation, Indoor Air Suppl. (14) (2004) S157–S167.

[6] S. Watanabe, T. Shimomura, H. Miyazaki, Thermal evaluation of a chair with fans as an individually controlled system, Build. Environ. 44 (2009) 1392–1398.

[7] S. Watanabe, A.K. Melikov, G.L. Knudsen, Design of an individually controlled system for an optimal thermal microenvironment, Build. Environ. 45 (2010) 549–558.

[8] J. Kaczmarczyk, A. Melikov, Z. Bolashikov, L. Nikolaev, P.O. Fanger, Human response to five designs of personalized ventilation, HVAC R. Res. 12 (2006) 367–384.

[9] S. Shahzad, J.K. Calautit, K. Calautit, B. Hughes, A.I. Aquino, Advanced personal comfort system (APCS) for the workplace: a review and case study, Energ. Build. 173 (2018) 689–709.

[10] Y. Ke, F. Wang, P. Xu, B. Yang, On the use of a novel nanoporous polyethylene (nanoPE) passive cooling material for personal thermal comfort management under uniform indoor environments, Build. Environ. 145 (2018) 85–95.

[11] Z. Ma, D. Zhao, F. Wang, R. Yang, CFD analysis of thermal comfort and energy saving of novel radiative cooling and heating textiles. Manuscript submitted for publication.

[12] W. Song, F. Wang, F. Wei, Hybrid cooling clothing to improve thermal comfort of office workers in a hot indoor environment, Build. Environ. 100 (2016) 92–101.

[13] Udayraj, Z. Li, Y. Ke, F. Wang, B. Yang, Personal cooling strategies to improve thermal comfort in warm indoor environments: comparison of a conventional desk fan and air ventilation clothing, Energ. Build. 174 (2018) 439–451.

[14] Z. Wang, K. Warren, M. Luo, X. He, H. Zhang, E. Arens, et al., Evaluating the comfort of thermally dynamic wearable devices, Build. Environ. 167 (2020) 106443.

[15] E. Arens, H. Zhang, C. Huizenga, Partial- and whole-body thermal sensation and comfort-Part I: uniform environmental conditions, J. Therm. Biol. 31 (2006) 53–59.

[16] N. Ghaddar, K. Ghali, S. Chehaitly, Assessing thermal comfort of active people in transitional spaces in presence of air movement, Energ. Build. 43 (2011) 2832–2842.

[17] J.D. Cotter, N.A.S. Taylor, The distribution of cutaneous sudomotor and alliesthesial thermosensitivity in mildly heat-stressed humans: an open-loop approach, J. Physiol. 565 (2005) 335–345.

[18] A.C. Boerstra, M. te Kulve, J. Toftum, M.G.L.C. Loomans, B. Olesen, J.L.M. Hensen, Comfort and performance impact of personal control over thermal environment in summer: results from a laboratory study, Build. Environ. 87 (2015) 315–326.

[19] F. Tartarini, S. Schiavon, T. Cheung, T. Hoyt, CBE Thermal Comfort Tool: online tool for thermal comfort calculations and visualizations, SoftwareX 12 (2020) 100563.

[20] American Society of Heating, Refrigerating and Air-Conditioning Engineers, in: Thermal Environmental Conditions for Human Occupancy, ASHRAE, Atlanta, GA, 2020 (ASHRAE Standard 55-2020).

[21] Y. Liu, M.H. Stowe, D. Bello, J. Sparer, R.J. Gore, M.R. Cullen, C.A. Redlich, S.R. Woskie, Skin exposure to aliphatic polyisocyanates in the auto body repair and refinishing industry: III. A personal exposure algorithm, Ann. Occup. Hyg. 53 (2009) 33–40.

[22] Z.J. Schlader, S.E. Simmons, S.R. Stannard, T. Mündel, The independent roles of temperature and thermal perception in the control of human thermoregulatory behavior, Physiol. Behav. 103 (2011) 217–224.

[23] C.J. Tyler, C. Sunderland, Cooling the neck region during exercise in the heat, J. Athl. Train. 46 (2011) 61–68.

[24] C.J. Stevens, A. Kittel, D.V. Sculley, R. Callister, L. Taylor, B.J. Dascombe, Running performance in the heat is improved by similar magnitude with pre-exercise cold-water immersion and mid-exercise face water spray, J. Sport Sci. 35 (2017) 798–805.

[25] Y. Cao, T.H. Lei, F. Wang, B. Yang, Head, facial and neck cooling as per-cooling modalities to improve exercise performance in the heat: a narrative review and practical applications, medRxiv, doi:10.1101/2021.05.31.21258125.

[26] C.J. Tyler, P. Wild, C. Sunderland, Practical neck cooling and time-trial running performance in a hot environment, Eur. J. Appl. Physiol. 110 (2010) 1063–1074.

[27] C. Sunderland, R. Stevens, B. Everson, C.J. Tyler, Neck-cooling improves repeated sprint performance in the heat, Front. Physiol. 6 (2015) 314.

[28] W. Pasut, E. Arens, H. Zhang, Y. Zhai, Enabling energy-efficient approaches to thermal comfort using room air motion, Build. Environ. 79 (2014) 13–19.

[29] S.H. Ho, L. Rosario, M.M. Rahman, Thermal comfort enhancement by using a ceiling fan, Appl. Therm. Eng. 29 (2009) 1648–1656.

[30] B. Yang, S. Schiavon, C. Sekhar, D. Cheong, K.W. Tham, W.W. Nazaroff, Cooling efficiency of a brushless direct current stand fan, Build. Environ. 85 (2015) 196–204.

[31] M. He, N. Li, Y. He, D. He, C. Song, The influence of personally controlled desk fan on comfort and energy consumption in hot and humid environments, Build. Environ. 123 (2017) 378–389.

[32] Y. He, N. Li, N. Li, J. Li, J. Yan, C. Tan, Control behaviors and thermal comfort in a shared room with desk fans and adjustable thermostat, Build. Environ. 136 (2018) 213–226.

[33] Y. Zhang, R. Zhao, Effect of local exposure on human responses, Build. Environ. 42 (2007) 2737–2745.

[34] A. Ghahramani, K, Zhang, K. Dutta, Z. Yang, B. Becerik-Gerber, Energy saving from temperature setpoints and deadband: quantifying the influence of building and system properties on savings, Appl. Energ. 165 (2016) 930–942.

[35] P.O. Fanger, Thermal environment-human requirements, Environmentalist 6 (1986) 275–278.

[36] M.L. Taub, Power to people: personal control in offices for thermal comfort and energy savings, Master Thesis, University of California, Berkeley, 2013.

[37] M. Luo, B. Cao, W. Ji, Q. Ouyang, B. Lin, Y. Zhu, The underlying linkage between personal control and thermal comfort: psychological or physical effects? Energ. Build. 111 (2016) 56–63.

[38] J. Kim, S. Schiavon, G. Brager, Personal comfort models-a new paradigm in thermal comfort for occupant-centric environmental control, Build. Environ. 132 (2018) 114–124.

[39] J. Maier, O. Zierke, H.J. Hoermann, P. Goerke, Effects of personal control for thermal comfort in long-distance trains, Energ. Build. 247 (2021) 111125.

[40] R. Rissetto, M. Schweiker, A. Wagner, Personalized ceiling fans: effects of air motion, air direction and personal control on thermal comfort, Energ. Build. 235 (2021) 110721.

